# Meningeal neutrophil infiltration drives inflammation-exacerbated pediatric stroke through IL-36γ signaling

**DOI:** 10.64898/2026.04.03.716365

**Authors:** Ching-Wen Chen, Hong-Ru Chen, Yi-Min Kuo, Alyson Prorock, Yongde Bao, Jonah Short-Miller, Melissa M. Kinkaid, Katia Sol-Church, Chia-Yi Kuan, Yu-Yo Sun

## Abstract

Infection and systemic inflammation have been identified as major risk factors for pediatric stroke; however, the immune mechanisms underlying these clinical observations remain poorly understood. Here, we utilized an LPS-sensitized photothrombotic stroke model (LPS/PT) to investigate the contribution of immune cell infiltration to pediatric stroke pathogenesis and identified a previously unrecognized route of neutrophil entry into the injured brain. Compared with photothrombotic stroke alone (PT), LPS/PT mice exhibited markedly increased neutrophil infiltration accompanied by a hyperactivated phenotype. Depletion of neutrophils, but not monocytes, significantly reduced infarct size in LPS/PT animals, indicating a central role for neutrophils in inflammation-exacerbated pediatric stroke. Notably, we observed that neutrophils first accumulated within the leptomeningeal space before entering the brain parenchyma during the early phase of stroke, suggesting that the meninges may serve as an initial staging site for neutrophil recruitment. Using KikGR photoconvertible reporter mice combined with two-photon imaging, we further demonstrated that neutrophils infiltrate the ischemic brain through a compromised meningeal barrier following stroke. Transcriptomic analysis of infiltrating neutrophils revealed distinct gene expression signatures between meningeal neutrophils and circulating blood-derived neutrophils. Among these, IL-36γ was highly enriched in meninges-associated neutrophils during pediatric stroke. Consistently, single-cell RNA sequencing of meningeal immune cells confirmed elevated IL-36γ expression within the neutrophil cluster in LPS/PT animals. Importantly, intracisternal administration of IL-36 receptor antagonist (IL-36Ra) or anti–ICAM-1 antibody significantly reduced infarct volume in this pediatric stroke model. Together, our findings identify the meningeal barrier as a critical gateway for neutrophil infiltration and reveal IL-36γ as a key inflammatory mediator regulating neuroinflammation in pediatric stroke, highlighting a potential therapeutic target for limiting immune-mediated brain injury.

## Introduction

Pediatric stroke is a rare but devastating neurological condition that can occur at any age during childhood and frequently results in lifelong neurological, behavioral, and socioeconomic consequences. Although the incidence of pediatric stroke is substantially lower than that in adults^1^, its long-term burden on patients and families remains profound. The etiology of pediatric stroke is complex and multifactorial, involving genetic predisposition, host susceptibility, and environmental triggers. Notably, epidemiological studies have consistently identified inflammation and infection—including maternal fever and systemic infection—as some of the strongest risk factors associated with pediatric stroke^2–5^. These observations suggest that inflammatory priming may critically influence stroke susceptibility and severity in the developing brain.

Despite these clinical observations, mechanistic understanding of how inflammatory signals exacerbate pediatric stroke remains limited, in part due to the lack of well-characterized preclinical models that recapitulate infection-sensitized pediatric stroke. Furthermore, the response of central nervous system (CNS) barrier systems in pediatric individuals following injury appears to be fundamentally different from that observed in adults. For example, studies in neonatal rodents have shown minimal Evans Blue leakage within the first 24 hours following transient middle cerebral artery occlusion (tMCAO)^6^, suggesting that blood–brain barrier (BBB) disruption is less pronounced in neonatal stroke compared with adult stroke models. Consistent with this observation, neutrophil infiltration into the ischemic brain is also markedly reduced in neonatal animals relative to adults^6^. These findings indicate that classical BBB breakdown may not be the dominant mechanism driving immune cell infiltration in the immature brain. In addition to differences in barrier responses, the functional properties of myeloid cells—including neutrophils and monocytes—also differ substantially between pediatric and adult immune systems. Pediatric myeloid cells often display reduced effector functions such as migration capacity and pathogen killing, yet exhibit increased plasticity compared with adult counterparts^7,8^. Our previous work further demonstrated that circulating pediatric monocytes can adopt microglia-like phenotypes following hypoxic-ischemic brain injury, highlighting the unique adaptability of developing immune cells within the CNS environment^9, 10^. However, how pediatric myeloid cells contribute to stroke pathogenesis and recovery remains poorly understood.

The brain is protected by multiple specialized barrier systems that regulate immune cell trafficking and molecular exchange between the CNS and the periphery. The best characterized of these is the BBB, which is formed by endothelial cells, astrocytic endfeet, and pericytes, and is reinforced by tight junction proteins that restrict paracellular permeability^11^. Beyond the BBB, additional barrier layers surrounding the CNS have recently gained increasing attention for their roles in neuroimmune regulation. One such barrier is the meninges barrier (glia limitans), a specialized astrocytic structure that forms a continuous barrier along the surface of the CNS parenchyma and around penetrating blood vessels. Astrocytic endfeet within the glia limitans attach to a basement membrane and can dynamically regulate immune cell access to the CNS through inducible junctional complexes^12,13^.

External to the glia limitans lies the pia mater, a delicate connective tissue layer composed primarily of fibroblast-like cells and collagen fibers that separates the cerebrospinal fluid (CSF) from the CNS parenchyma. Numerous blood vessels traverse the pia mater before penetrating the brain, making this interface a potential site of immune surveillance. Above the pia mater lies the arachnoid barrier layer, which consists of specialized epithelial-like cells expressing tight junction proteins and adhesion molecules such as E-cadherin and VE-cadherin^14,15^. These arachnoid barrier cells form the boundary of the subarachnoid space (SAS) and play a key role in restricting the entry of immune mediators and cells into the CNS during both homeostasis and inflammation. Beneath the skull, the dura mater represents another immunologically active compartment enriched with circulating and tissue-resident immune cells. The recent discovery of functional meningeal lymphatic vessels within the dura mater has further highlighted the importance of meningeal structures in coordinating immune–CNS interactions^16^. Increasing evidence suggests that immune cells within the meninges can serve as early responders to CNS injury and may influence both neuroinflammation and neurological function^17,18^.

Interleukin-36 (IL-36) is a potent pro-inflammatory cytokine belonging to the IL-1 cytokine family that functions as an important alarmin at epithelial and mucosal barrier surfaces. The IL-36 cytokine family consists of three agonistic ligands—IL-36α, IL-36β, and IL-36γ—as well as a natural antagonist, IL-36 receptor antagonist (IL-36Ra), which competitively inhibits IL-36 receptor signaling^19^. IL-36 agonists are secreted as inactive precursors that require N-terminal proteolytic processing by neutrophil-derived proteases such as elastase and cathepsin G to achieve full biological activity^20^. Once activated, IL-36 signaling plays a critical role in regulating barrier integrity, promoting epithelial repair, modulating tight junction permeability, and initiating robust antimicrobial responses. IL-36 signaling can also drive the IL-23–IL-17–IL-22 cytokine axis, leading to the activation and expansion of Th17 cells, γδ T cells, and type 3 innate lymphoid cells (ILC3s)^21,22^. Importantly, IL-36 stimulates epithelial cells to produce potent neutrophil-attracting chemokines such as CXCL1 and CXCL8 (IL-8), thereby amplifying neutrophil recruitment to inflamed barrier tissues^23^. Recruited neutrophils subsequently release additional proteases that convert inactive IL-36 into its fully active form, establishing a feed-forward inflammatory amplification loop at barrier sites.

In the present study, we utilized a pediatric stroke model in which animals were pre-sensitized with a low dose of lipopolysaccharide (LPS) prior to ischemic injury. We found that systemic inflammatory priming markedly exacerbated stroke severity in a neutrophil-dependent manner, whereas monocyte depletion did not significantly alter disease outcomes. Using flow cytometry analysis of immune cell infiltration, we observed that neutrophils first accumulated within the meninges before entering the ischemic brain parenchyma. Further imaging studies using two-photon microscopy and photoconvertible KikGR reporter mice demonstrated that early meningeal barrier disruption provided a potential route for neutrophil migration into the injured brain. Transcriptomic analysis, including bulk RNA sequencing and single-cell RNA sequencing, revealed distinct transcriptional profiles between blood-brone neutrophils and meninges-infiltrating neutrophils, with a striking enrichment of IL-36γ expression in the meningeal neutrophil population. Finally, we show that intracisternal administration of anti-ICAM-1 antibody to inhibit neutrophil migration, as well as IL-36 receptor antagonist (IL-36Ra) to block IL-36 signaling, significantly reduced infarct volume in this pediatric stroke model. Together, our findings reveal a previously unrecognized meningeal route for immune cell infiltration in pediatric stroke and identify IL-36γ as a key inflammatory mediator regulating meningeal barrier dysfunction during CNS injury.

## Materials and Methods

### Animals

C57BL/6J, CCR2^RFP/RFP^ (JAX#017586), and GFAP-Cre (JAX#012886) mice were purchased from The Jackson Laboratory. KikGR photoconvertible mice^24^ were kindly provided by Dr. Jonathan Kipnis. All mice were maintained in ventilated cages under specific pathogen–free conditions with standard laboratory chow and water provided ad libitum. All experimental procedures were approved by the Institutional Animal Care and Use Committee (IACUC) at the University of Virginia and conducted in accordance with the guidelines of the Association for Assessment and Accreditation of Laboratory Animal Care (AAALAC).

### Pediatric photothrombotic middle cerebral artery occlusion (MCAO) model

Postnatal day 16 (P16) mice of both sexes were used to model pediatric stroke. For inflammation-sensitized stroke experiments, mice received a single intraperitoneal injection of lipopolysaccharide (LPS; 0.3 mg/kg in 50 µL PBS) one hour prior to photothrombotic MCAO surgery^25^. Briefly, mice were anesthetized with isoflurane, and the left scalp was surgically opened to expose the proximal segment of the left middle cerebral artery (MCA). Rose Bengal (Thermo Fisher Scientific) dissolved in saline was administered via retro-orbital injection (25 mg/kg in 50 µL). A green laser (543.5 nm, 5 mW; Melles Griot, Carlsbad, CA) was then directed at the proximal MCA for 15 minutes to induce photothrombotic occlusion. Following surgery, the scalp was sutured and mice were allowed to recover on a heating pad.

For infarct volume analysis, brains were harvested and sectioned into 1-mm-thick coronal slices. Brain sections were incubated in 2,3,5-triphenyltetrazolium chloride (TTC) solution for 20 minutes at room temperature. TTC-negative areas representing infarcted tissue were quantified using ImageJ software, and infarct volume was calculated as the sum of infarct areas across all sections. For neutrophil depletion experiments, mice received intraperitoneal injections of anti-Ly6G antibody (1A8, BioXCell; 100 µg per dose) at P15, P16, and P17.

### Intra-cisterna magna (ICM) injection

For intra-cisterna magna (ICM) injection^26^, P16 mice were anesthetized with ketamine. A total volume of 2 µL was injected into the cisterna magna using a Nanoliter 2020 injector at an infusion rate of 3 nL/sec. After injection, the needle was left in place for an additional 5 minutes to allow diffusion of the injected solution before withdrawal. The following reagents were used for ICM administration: Fluorescein isothiocyanate (FITC)–dextran (70 kDa; 10 mg/mL; Sigma-Aldrich); Alexa Fluor 488–conjugated bovine serum albumin (BSA; 10 mg/mL; Invitrogen); Anti-CD45.2 antibody (clone 104; 0.5 mg/mL; BioLegend); Anti-ICAM-1 antibody (5 mg/mL; BioXCell); Recombinant mouse IL-36 receptor antagonist (IL-36Ra; 250 µg/mL; R&D Systems)

### Flow cytometry

Mice were transcardially perfused with phosphate-buffered saline (PBS) to remove circulating blood cells prior to tissue collection. Ipsilateral and contralateral brain hemispheres and skull-associated meninges (dura) were harvested at designated time points following stroke^26^. Brain hemispheres were mechanically dissociated, and immune cells were isolated using discontinuous Percoll gradients (GE Healthcare). For meningeal immune cell isolation, dura mater was carefully dissected from the skull and enzymatically digested for 25 minutes at 37°C in RPMI containing: Collagenase VIII (1.4 U/mL; Sigma-Aldrich), Collagenase D (1 mg/mL; Roche) and DNase I (35 U/mL; Sigma-Aldrich). Following digestion, cells were washed and subjected to flow cytometry staining.

During staining, cell suspensions were stained with LIVE/DEAD Fixable Aqua Dead Cell Stain (Thermo Fisher Scientific) along with fluorophore-conjugated antibodies including: CD45 (30-F11; BioLegend), CD11b (M1/70; BioLegend), Ly6G (1A8; BioLegend), Ly6C (HK1.4; BioLegend), CD62L (MEL-14; BioLegend), CXCR2 (SA044G4; BioLegend), ICAM-1 (YN1/1.7.4; BioLegend), iNOS (W16030C; BioLegend). Absolute immune cell numbers were determined using CountBright™ absolute counting beads (Thermo Fisher Scientific). Samples were acquired on an Attune flow cytometer (Applied Biosystems) and analyzed using FlowJo v10 software.

### Immunohistochemistry

Brains were harvested at designated time points and fixed overnight in 4% paraformaldehyde (PFA) at 4°C. Fixed brains were cryoprotected sequentially in 15% and 30% sucrose, embedded, and stored at −80°C until sectioning. Coronal cryosections (20 µm) were prepared and permeabilized with 1% Triton X-100 in PBS for 10 minutes, followed by blocking in 3% normal goat serum (NGS) for 1 hour. Sections were incubated overnight with primary antibodies diluted in 1% NGS and 0.1% Triton X-100 in PBS. After washing, sections were incubated with corresponding secondary antibodies for 3 hours at room temperature. For meningeal staining, skulls containing dura mater were fixed in 2% PFA for 2 hours at 4°C. Dura mater was carefully dissected and stained using free-floating protocols similar to those used for brain sections. Primary antibodies included: Isolectin GS-IB4 (Invitrogen™), Ly6G (1A8; AF488 or AF647 conjugated; BioLegend), Laminin (ab11575; Abcam), GFAP (ab4674; Abcam), IL-36γ (United States Biological; 140948)

### Meningeal barrier permeability assay

To assess meningeal barrier permeability, LPS-sensitized stroke mice were injected via ICM with 2 µL FITC–dextran (70 kDa; 10 mg/mL) 30 minutes prior to sacrifice. Brains were fixed in 4% PFA overnight, and 100 µm free-floating brain sections were stained with anti-Ly6G antibody. For quantitative analysis of meningeal barrier leakage, Alexa Fluor 488–conjugated bovine serum albumin (BSA-AF488) was injected via ICM at various time points following stroke. Mice were sacrificed 1 hour after injection. Ipsilateral cortical tissues were collected, weighed, and homogenized. Total protein extracts were prepared and BSA-AF488 fluorescence was quantified using a microplate reader (SpectraMax M3, Molecular Devices).

### Photoconversion of KikGR mice

P16 KikGR mice underwent LPS-sensitized photothrombotic MCAO as described above. At 6 hours post-MCAO, mice were anesthetized and the scalp was reopened to expose the ipsilateral skull. Photoconversion was performed using a violet optogenetic LED module (Prizmatix) for 30 seconds per spot at intensity level 6^26^. Five spots across the ipsilateral skull were illuminated with the light source positioned approximately 1 cm above the skull surface. Following photoconversion, mice were allowed to recover on a heating pad. Mice were sacrificed at 9 and 12 hours post-stroke (corresponding to 3 and 6 hours after photoconversion). KikR-positive neutrophils in blood, meninges, and brain tissue were quantified by flow cytometry.

### Intravital confocal imaging

Neutrophils were isolated from the bone marrow of P16 mice using a Neutrophil Isolation Kit (Miltenyi Biotec). Purified neutrophils were labeled ex vivo with CellTracker Orange CMTMR dye (Thermo Fisher Scientific) for 30 minutes following the manufacturer’s protocol. A total of 5 × 10^6^ labeled neutrophils were adoptively transferred via retro-orbital injection into GFAP-Cre × ZsGreen reporter mice. After a 1-hour resting period, mice were treated with LPS and subjected to photothrombotic MCAO. Three hours after stroke induction, a cranial window was installed over the ipsilateral parietal bone to allow intravital imaging of neutrophil migration between the meninges and brain parenchyma. Two-photon imaging data were processed and analyzed using Imaris software.

### Bulk RNA sequencing and quantitative real-time PCR

To distinguish neutrophils entering the brain via the meningeal route from circulating neutrophils, mice received ICM injection of Alexa Fluor 700–conjugated anti-CD45.2 antibody (1 µg; BioLegend) and intravenous injection of Alexa Fluor 488–conjugated anti-CD45 antibody (5 µg; BioLegend) at 23 hours post LPS-sensitized stroke. After one hour, mice were sacrificed and immune cells were isolated from blood, meninges, and ipsilateral brain parenchyma. Cells were stained with LIVE/DEAD Fixable Aqua dye and antibodies against CD11b and Ly6G, followed by fluorescence-activated cell sorting (FACS) using a BD Influx Cell Sorter at the University of Virginia Flow Cytometry Core Facility. Three to five mice were pooled per biological sample, yielding approximately 5,000–20,000 neutrophils per sample. RNA was extracted and quality assessed using High Sensitivity RNA ScreenTape. Reverse transcription, cDNA amplification, and indexed libraries were generated using the NEBNext Ultra II Directional RNA Library Prep Kit. Pooled libraries were sequenced on the Illumina NextSeq2000 platform with 1 × 100 bp single-end reads by the University of Virginia Genome Analysis and Technology Core (RRID:SCR_018883), and the demultiplexed QC reads shared for downstream analysis. For validation experiments, neutrophils were independently isolated from five biological replicates and RNA was extracted using TRIzol reagent (Invitrogen). Quantitative real-time PCR was performed using a Bio-Rad CFX96 system with SYBR Green master mix. The following IL-36γ primers were used: Forward: AGAGTAACCCCAGTCAGCGTG Reverse: AGGGTGGTGGTACAAATCCAA

### Single-cell RNA sequencing

Meningeal tissues were collected from LPS-sensitized stroke mice and sham controls at 24 hours post-stroke. Dura mater was dissected and enzymatically digested as described above. Cells were stained with LIVE/DEAD Fixable Aqua dye and CD45 antibody, and CD45+ immune cells were sorted by flow cytometry.

Single-cell libraries were generated using the Chromium Next GEM Single Cell 3′ v3.1 platform (10x Genomics) according to the manufacturer’s instructions. Approximately 7,000–8,000 CD45+ cells from each condition were loaded into the Chromium controller for droplet-based barcoding and library preparation. Libraries were sequenced on an Illumina NextSeq 2000 platform and processed at the Genome Analysis and Technology Core at the University of Virginia.

## Results

### LPS priming amplifies neutrophil-dominant inflammation and increases infarct burden in a pediatric stroke model

Postnatal day 16 (P16) mice are approximately similar to late infancy in humans^27^, which is a developmental window associated with high incidence of pediatric arterial ischemic stroke. To model the clinical association between infection/inflammation and pediatric stroke risk, we combined low-dose LPS priming (0.3 mg/kg) with photothrombotic ischemia (LPS/PT). Compared with photothrombosis alone (PT), LPS/PT animals developed significantly larger infarcts at day 3 post-injury (Fig. 1a), establishing that systemic inflammatory priming markedly exacerbates ischemic tissue damage in the immature brain. To define the inflammatory milieu underlying this sensitization, we quantified cytokine and chemokine mRNA in the ipsilateral hemisphere across four conditions (untreated, LPS only, PT only, and LPS/PT) at early time points. LPS/PT induced robust increases in pro-inflammatory cytokines (Il1a, Il1b) and neutrophil-associated chemokines (Cxcl1, Cxcl2) that were detectable as early as 4 h post-stroke (Fig. 1b), promoting an environment to rapid neutrophil recruitment. We next profiled leukocyte influx by flow cytometry at 24 h and 72 h post-stroke (gating strategy in Fig. 1c). Both Ly6Chi monocytes (CD45^hi^Ly6G^−^Ly6C^hi^) and neutrophils (CD45^hi^Ly6G^+^) increased at 24 h and declined by 72 h; however, LPS priming produced a prominent shift toward a neutrophil-dominant infiltrate. Specifically, neutrophil numbers were significantly higher in LPS/PT than in PT at 24 h (Fig. 1d–e), whereas monocyte accumulation was not increased by LPS priming and represented a smaller fraction of the infiltrate in LPS/PT (Fig. 1d). Immunohistochemistry further validated parenchymal entry of neutrophils. Ly6G+ cells were detected outside IB4+ vascular structures in the ipsilateral cortex, indicating extravasation margination into tissue rather than intravascular (Fig. 1f). Notably, neutrophils were also abundant along leptomeningeal interfaces, including the pial surface and perivascular spaces (PVS), suggesting that brain surface compartments may represent early sites of neutrophil positioning during pediatric ischemic injury (Fig. 1f).

**Figure 1.**
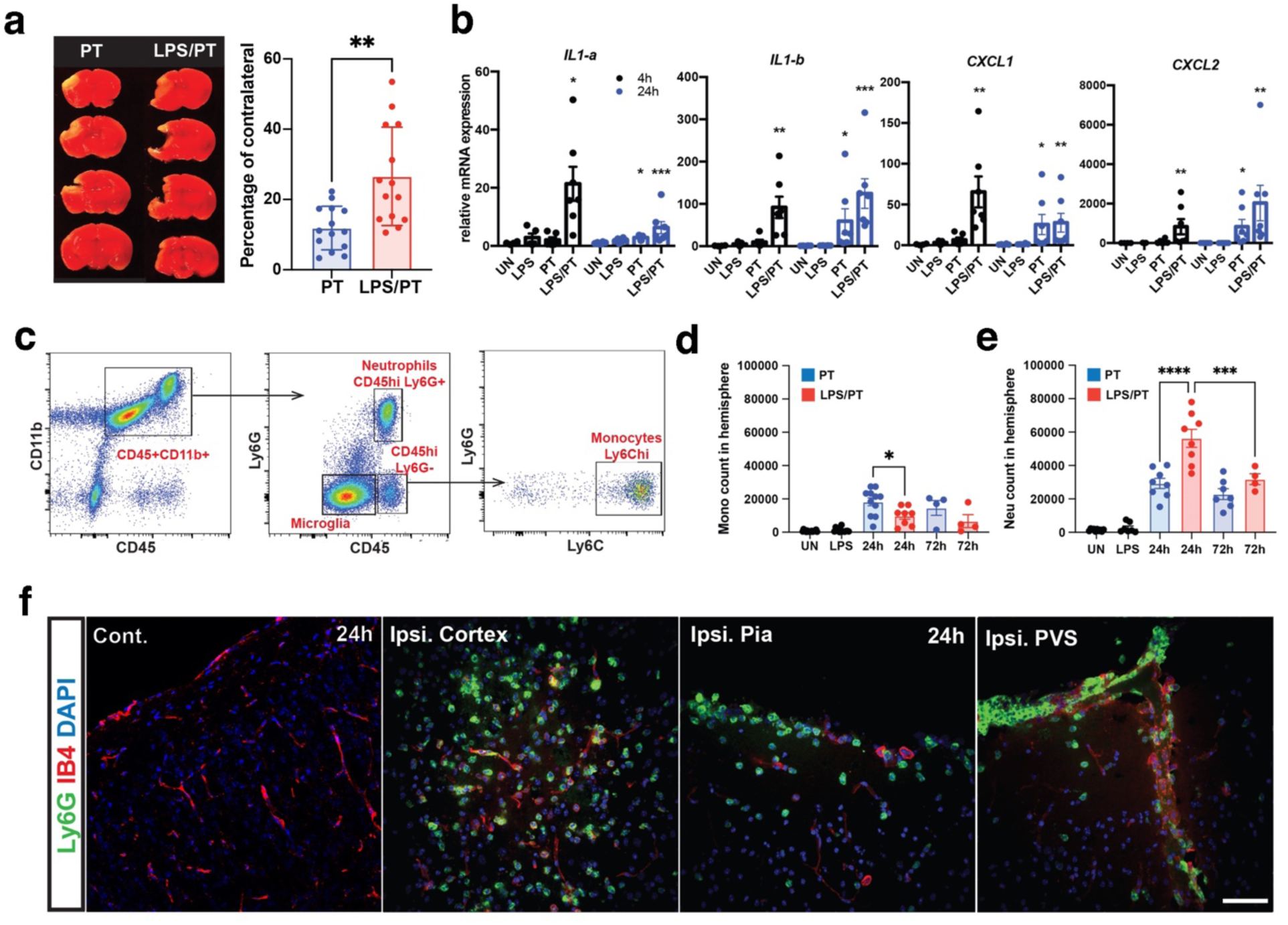
LPS sensitization promotes accumulation of hyperactivated neutrophils in the ipsilateral brain parenchyma after pediatric stroke. **a**, Postnatal day 16 (P16) mice were randomly assigned to two experimental groups. One group underwent photothrombotic (PT) occlusion of a middle cerebral artery (MCA) branch to induce pediatric ischemic stroke (PT, n=14). The second group received intraperitoneal lipopolysaccharide (LPS; 0.3 mg/kg, low-dose sensitization) 1 h prior to PT surgery (LPS/PT, n=14). Animals were euthanized 3 days post-stroke, and brain sections were stained with 2,3,5-triphenyltetrazolium chloride (TTC). Representative TTC-stained sections are shown, and infarct size were quantified as a percentage of the contralateral hemisphere. Data were presented as mean ± S.E.M. **p < 0.01 compared with the contralateral hemisphere by unpaired t-test. **b**, Animals from each experimental group were euthanized at 4 h or 24 h post-surgery, and ipsilateral brain hemispheres were collected for quantitative PCR analysis of Il1a, Il1b, Cxcl1, and Cxcl2 mRNA expression. Data were pooled from two independent experiments (n = 6 per group) and analyzed using two-way ANOVA followed by Tukey’s post hoc test. *p < 0.05, **p < 0.01, ***p < 0.001, compared with the uninjured (UN) group. **c-e**, Animals were euthanized 24 h after treatment, and ipsilateral brain hemispheres were harvested for flow cytometric analysis. **c**, Flow cytometric gating strategy for analysis of immune cell populations in the brain. After singlet and live cell gating, CD45⁺CD11b⁺ cells were selected. Neutrophils were identified as CD45hiLy6G⁺ cells, and monocytes were identified as CD45hiLy6G⁻Ly6Chi cells. Absolute numbers of infiltrating monocytes (Mono, **d**) and neutrophils (Neu, **e**) are shown. Data were analyzed by one-way ANOVA with Tukey’s post hoc test. *p < 0.05, ***p < 0.001, ****p < 0.0001 compared with the UN group. N > 6 per group. **f**, High-magnification confocal images demonstrating neutrophil accumulation in the ipsilateral cerebral cortex, including the pial surface (Pia), perivascular spaces (PVS), and deep cortical regions, 24 h after LPS/PT treatment. Neutrophils were identified by anti-Ly6G immunostaining, and blood vessels were labeled with isolectin B4 (IB4). The contralateral hemisphere served as an internal control. Representative images are shown. Scale bar: 50 mm. N=3 for each tissue sample.

Because inflammatory priming can alter neutrophil functional states, we assessed activation-associated markers on tissue-infiltrating neutrophils. Neutrophils recovered from LPS/PT ipsilateral hemispheres displayed increased ICAM-1 expression and elevated iNOS, consistent with a pro-inflammatory activation program^28^ (Fig. 2a). To determine whether neutrophils drive the increased infarct burden in LPS/PT, we depleted neutrophils using anti-Ly6G (1A8) and confirmed efficient depletion in peripheral blood (Sup. Fig. 1a). Neutrophil depletion significantly reduced infarct volume in LPS/PT animals, whereas it did not alter infarct size in PT animals (Fig. 2b-c). Next, we took advantage of genetic ablation of CCR2 in CCR2^RFP/RFP^ animals to suppress monocyte infiltrations after stroke. In contrast, genetic suppression of CCR2-dependent monocyte recruitment (Ccr2 ^RFP/RFP^) did not reduce infarct size in LPS/PT animals compared with CCR2-intact littermate controls (Ccr2 ^RFP/-^). Together, these data identify neutrophils as the principal driver of inflammation-sensitized pediatric stroke pathology in this model.

**Figure 2.**
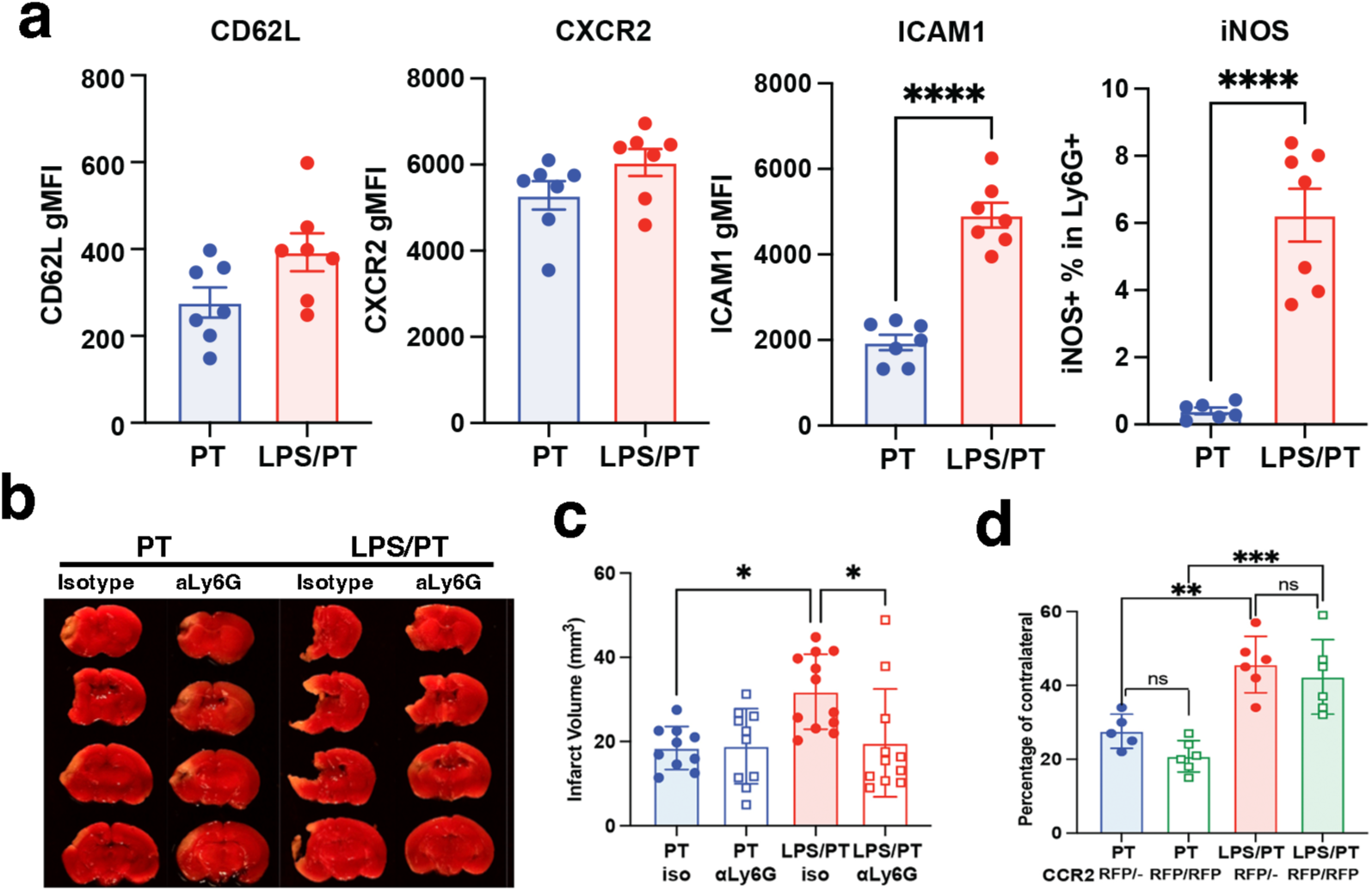
Neutrophil depletion, but not CCR2 deficiency, attenuates infarct size in LPS-sensitized pediatric stroke. **a,** Expression of neutrophil activation markers (ICAM1, iNOS, CD62L and CXCR2) in the ipsilateral hemisphere was assessed by flow cytometry at 24 h post-stroke in PT and LPS/PT groups (N = 7 per group). Data were presented as mean ± S.E.M. ****p < 0.0001 compared with the PT group by unpaired t-test. **b-c,** Mice were treated with a neutrophil-depleting anti-Ly6G antibody (clone 1A8) or an isotype control antibody beginning one day prior to stroke induction and continuing daily for three consecutive days (P15–P17). Brain sections (1mm sections) were collected 3 days post-stroke and stained with TTC. Infarct volumes were quantified in mm3. (N> 10 per group). *p < 0.05 by one-way ANOVA. **d,** CCR2–RFP knock-in mice, including homozygous CCR2-deficient (CCR2RFP/RFP) and heterozygous CCR2-sufficient (CCR2RFP/+) littermates, were subjected to PT or LPS/PT. Brain sections were harvested 3 days post-stroke and infarct volumes were quantified by TTC staining and expressed as a percentage of the contralateral hemisphere (N > 5 per group). ns, Not significant, p>0.05; **p < 0.01; ***p < 0.001 by one-way ANOVA.

### Neutrophils preferentially accumulate within meningeal compartments before entering ischemic parenchyma at early time point of stroke

Given the dominant contribution of neutrophils to injury in LPS/PT, we next sought to define the route(s) of neutrophil entry into the brain. In adult stroke, BBB disruption and endothelial activation can occur rapidly after ischemia^29^; however, in our pediatric LPS/PT model we did not detect appreciable neutrophil accumulation within the injured parenchyma at 6h or even 12h post-stroke (Fig. 3a), despite early induction of neutrophil chemoattractants (Fig. 1b). This temporal dissociation suggested that neutrophils may initially localize to non-parenchymal compartments. Consistent with this hypothesis, confocal imaging revealed prominent neutrophil accumulation at the pial surface at 24 h post-LPS/PT (Sup. Fig. 1b). We therefore quantified immune cells in parenchyma and meninges dura across early time points. Flow cytometric analysis demonstrated that neutrophils in the ipsilateral parenchyma increased only modestly until 12 h and rose markedly by 24 h post-LPS/PT (Fig. 3a). In striking contrast, neutrophils accumulated in meningeal dura compartments as early as 6 h and remained elevated through 24 h (Fig. 3b). Whole-mount meningeal preparations further revealed neutrophil enrichment on the ipsilateral dura, including regions adjacent to the superior sagittal and transverse sinuses (Fig. 3c). Importantly, compared with PT alone, LPS/PT selectively increased meningeal neutrophil numbers without a corresponding increase in meningeal monocytes (Sup. Fig. 1c), indicating that inflammatory priming preferentially amplifies neutrophil recruitment to meninges in this pediatric stroke context. High-magnification imaging of meningeal vessels demonstrated Ly6G+ cells located outside vascular lumens, consistent with active transendothelial migration into meningeal tissues/leptomeningeal spaces (Fig. 3d). Because neutrophil entry directly across the brain–pia interface has not been established in stroke, we next mapped neutrophil positions relative to anatomical barrier layers. We delineated the pia/basement membrane with laminin and the glia limitans with GFAP. At 6 h post-LPS/PT, laminin staining was preserved and the glia limitans remained intact; at this time point, neutrophils were primarily positioned on the CSF-facing side of the pia (Fig. 3e). By 12 h post-LPS/PT, neutrophils were observed traversing the pia and glia limitans (Fig. 3e, yellow arrows). By 24 h post-LPS/PT, laminin signal was markedly reduced and the glia limitans appeared disrupted, coinciding with substantially increased parenchymal neutrophil accumulation (Fig. 3e). These findings indicate that neutrophils accumulate early at meningeal interfaces and subsequently penetrate the brain surface barrier coincident with disruption of the pia–glia limitans boundary.

**Figure 3.**
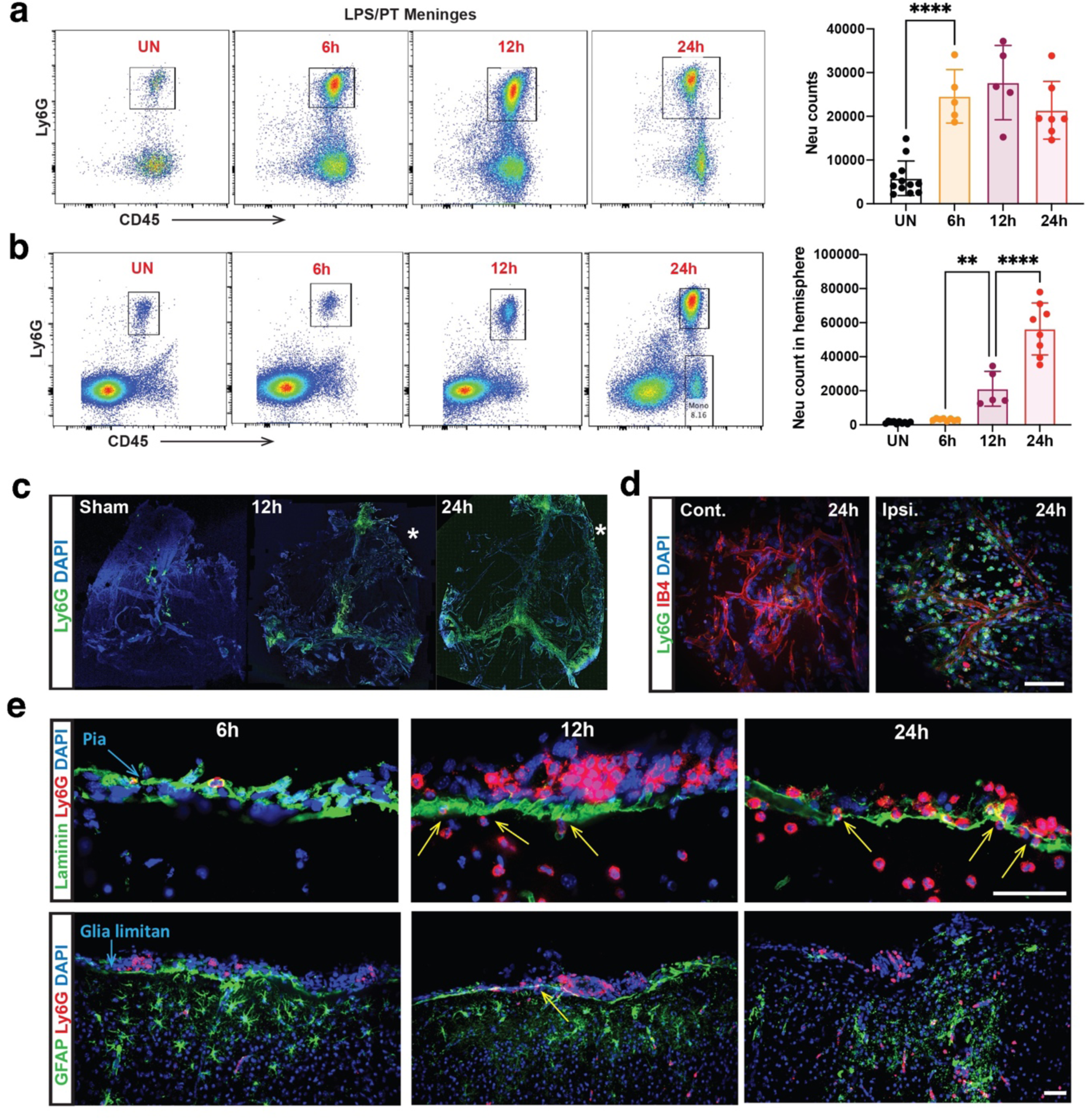
Neutrophils accumulate at the leptomeninges prior to parenchymal infiltration during the early phase of LPS-sensitized pediatric stroke. **a-b,** Neutrophil numbers in the ipsilateral brain hemisphere (A) and meninges (dura mater, B) were quantified by flow cytometry at 6 h, 12 h, and 24 h following LPS/PT. Representative flow cytometric plots and corresponding quantitative analyses are shown. Data were presented as mean ± S.E.M (N> 5 per group). **p < 0.01; ****p < 0.0001 by one-way ANOVA. **c,** Whole-mount preparations of the meninges (dura mater) were harvested at 12 h and 24 h, and immunostained with anti-Ly6G to visualize neutrophils. The asterisk denotes the site of photothrombotic injury. Representative images from each experimental group are shown (N=3 per group). **d,** High-magnification confocal images of the contralateral and ipsilateral meninges at 24 h post-LPS/PT, stained with anti-Ly6G to label neutrophils and isolectin B4 (IB4) to visualize the vasculature. Representative images are shown (N=3 per group). **e,** Immunostaining for laminin and GFAP was performed to delineate the pia mater and glia limitans at the cortical surface. Neutrophils were identified by Ly6G staining. Representative images from 6 h, 12 h, and 24 h post-LPS/PT are shown (N=3 per group).

### The brain–meningeal barrier becomes permeable early after LPS/PT, temporally coinciding with neutrophil transmigration

We next tested whether stroke triggers functional impairment of the brain–meningeal barrier in LPS/PT animals. We injected FITC–dextran into the cisterna magna and examined tracer distribution in the ipsilateral hemisphere 30 min later. At 24 h post-LPS/PT, FITC–dextran distributed from CSF into the ipsilateral parenchyma via leptomeningeal interfaces, in regions where neutrophils were concurrently detected (Fig. 4a), supporting the presence of meningeal barrier leakage. To quantify the onset of barrier permeability, we adapted a tracer-based leakage assay used to measure BBB disruption^29^ and injected Alexa Fluor 488–conjugated BSA into the subarachnoid space via intracisterna magna (ICM) administration at defined time points after LPS/PT. One hour after ICM injection, fluorescence intensity in the ipsilateral cortex was measured. BSA accumulation was significantly increased by 12 h post-LPS/PT (Fig. 4b), indicating that brain–meningeal barrier permeability emerges early after injury. Notably, this timing aligns with the appearance of neutrophils traversing the pia and glia limitans (Fig. 3e), supporting a model in which early meningeal barrier impairment provides an access route for neutrophils into injured cortex.

**Figure 4.**
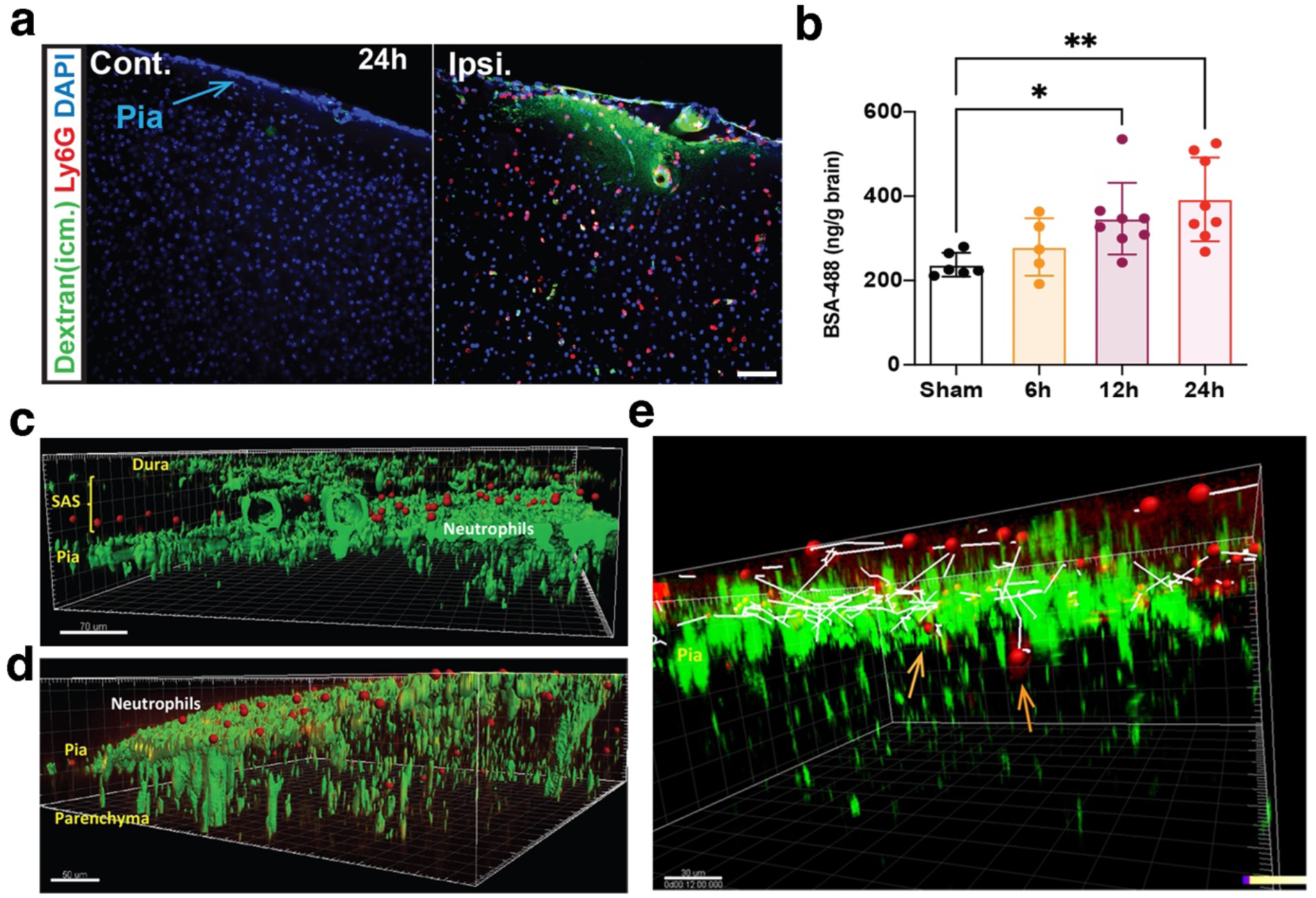
Impairment of the brain–meningeal barrier during LPS-sensitized pediatric stroke facilitates neutrophil migration across the pia mater. **a,** To assess brain–meningeal barrier integrity, fluorescein isothiocyanate (FITC)–dextran (70 kDa; 10 mg/mL; 2 µL) was injected into the cisterna magna (i.c.m.) at 24 h post-stroke. Animals were euthanized 1 h after intracisternal injection, and brain sections were analyzed by confocal microscopy to visualize FITC–dextran leakage into the cortical parenchyma. Neutrophils were identified by anti-Ly6G immunostaining. Representative images are shown (N=3 per group). **b,** Brain–meningeal barrier permeability was quantitatively assessed at indicated time points after stroke at 6 h, 12 h or 24 h by i.c.m. injection of Alexa Fluor 488–conjugated bovine serum albumin (BSA-488; 10 mg/mL; 2 µL). Animals were euthanized 1 h post-injection, and ipsilateral cortical tissue was homogenized for total protein extraction. BSA-488 fluorescence intensity was measured to determine barrier permeability. Data are presented as mean ± S.E.M. (n > 5 per group). *p < 0.05; **p < 0.01 compared with the sham group by one-way ANOVA. **c-e,** Two-photon intravital imaging was performed to visualize neutrophil localization and migration across the pia mater and glia limitans following stroke. Neutrophils were isolated from P16 mouse bone marrow using a neutrophil isolation kit and labeled ex vivo with CellTracker™ Orange CMTMR dye. Labeled neutrophils (5 × 10⁶ cells) were adoptively transferred into GFAP-Cre × ZsGreen (GFAP-GFP) reporter mice. Recipient mice were sensitized with LPS and subjected to photothrombotic stroke as described above. A cranial window was established over the ipsilateral parietal bone at 3 h post-LPS/PT, and two-photon imaging was performed approximately 16 h post-LPS/PT (n=4). Imaging of the dura mater (Dura) and subarachnoid space (SAS), with neutrophils visualized as red fluorescent cells (**c**). Imaging of the pia mater (Pia) and cortical parenchyma (**d**). Representative time-lapse tracking of neutrophil migration (white trajectory) over a 30-min imaging period (**e**).

### Intravital imaging and photo-converted tracing reveal neutrophil migration from meningeal compartments into injured cortex via a disrupted pia–glia limitans interface

Although our histological and flow cytometric analyses suggested that neutrophils accumulate in leptomeningeal spaces and traverse brain surface barriers, direct visualization of this route has not been reported in stroke. We therefore employed in vivo two-photon microscopy to directly monitor neutrophil localization relative to the glia limitans in live animals. To visualize glial boundary structures, we generated GFAP-GFP reporter mice (GFAP-Cre × ZsGreen), which displayed strong fluorescence along the glia limitans layer. Notably, additional fluorescence signal near vascular structures is also observed, consistent with reported GFAP expression in certain perivascular fibroblast-like populations^30^ (Fig 4c). We then isolated bone marrow neutrophils from donor P16 animals, labeled them with CMTMR, and adoptively transferred them into recipient GFAP-GFP mice prior to LPS/PT. At 16h post-LPS/PT, labeled neutrophils were readily detected in the subarachnoid space and along the pial surface (Fig. 4c), consistent with early leptomeningeal positioning. We also observed neutrophils beneath the glia limitans within the parenchyma (Fig. 4d), in agreement with our fixed-tissue analyses (Fig. 3e). Time-lapse imaging captured individual neutrophils actively transmigrating from the CSF-facing side of the pia into the parenchyma (Fig. 4e; Sup. Video 1), providing direct evidence that neutrophils can enter injured pediatric cortex via a meningeal route during stroke.

To independently validate the origin of early infiltrating neutrophils, we used Kikume Green-Red (KikGR) photoconvertible reporter mice^24^. Following LPS/PT, we photoconverted meningeal immune cells using a violet laser at 6h post-injury, a time when meningeal neutrophils were abundant (Fig. 3b, Fig 5a). Photoconversion shifted approximately ∼15% of meningeal neutrophils from KikG^+^KikR^−^ to KikG^−^KikR^+^ (Fig. 5b). Importantly, KikG–KikR+ neutrophils were not detected in peripheral blood after meningeal photoconversion (Fig. 5c), supporting spatial confinement of converted cells. We then assessed the ipsilateral hemisphere for photoconverted KikG^−^KikR^+^ neutrophils. At 3 h post-photoconversion (9 h post-LPS/PT), approximately ∼18% of parenchymal infiltrating neutrophils were KikG^−^KikR^+^, indicating recent migration from the photoconverted meningeal compartment. At 6 h post-photoconversion (12 h post-LPS/PT), the absolute number of photoconverted neutrophils increased while their fraction decreased, resulting from continued meningeal contribution and increasing influx of later-arriving blood-borne neutrophils over time. Together, intravital imaging and photo-converted tracing establish that meningeal compartments provide a functional and temporally early route for neutrophil entry into the injured pediatric brain.

**Figure 5.**
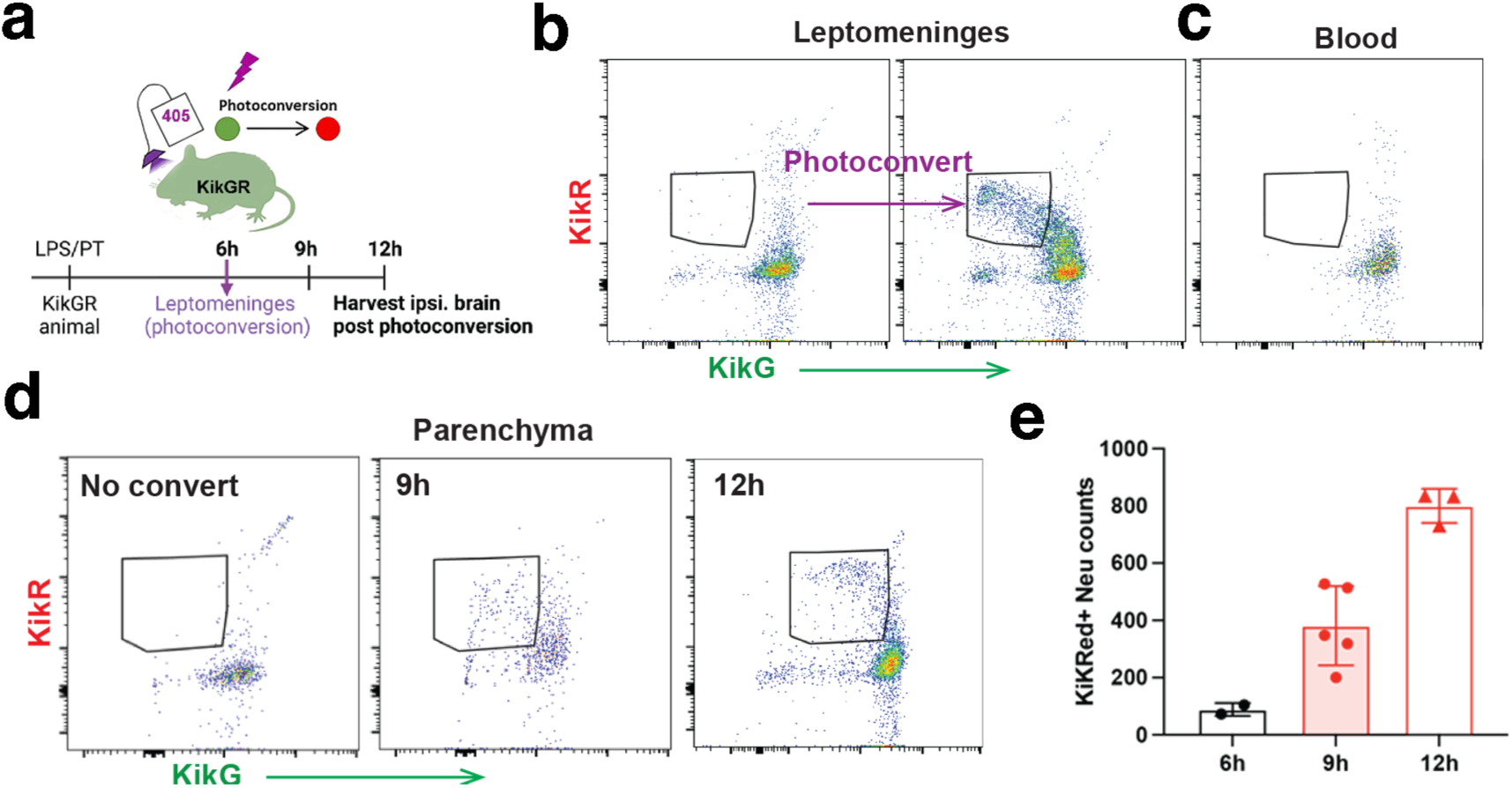
Photoconverted neutrophils migrate from the leptomeningeal space into the brain parenchyma following LPS-sensitized pediatric stroke. **a,** Schematic illustrating the experimental timeline for in-vivo photoconversion in KikGR reporter mice. At 6 h post-stroke, corresponding to the peak of neutrophil accumulation in the leptomeninges, a violet laser was applied to multiple regions of the intact skull to selectively photoconvert cells localized within the leptomeningeal space. Animals were euthanized at 6 h, 9 h, or 12 h after LPS/PT for analysis. **b-d,** Representative flow cytometric analyses demonstrating photoconverted neutrophils, identified by the transition from KikGreen⁺KikRed⁻ to KikGreen⁻KikRed⁺ within CD45hiCD11bhiLy6G⁺ cells. Detection of photoconverted neutrophils in the leptomeninges following localized skull illumination (**b**). Peripheral blood analysis showing absence of KikRed⁺ neutrophils, confirming spatially restricted photoconversion (**c**). Representative flow cytometric plots of photoconverted neutrophils identified in the ipsilateral brain parenchyma at 6 h, 9 h, and 12 h post-LPS/PT (**d**). Quantification of KikRed⁺ neutrophils (CD45hiCD11bhiLy6G⁺) in the ipsilateral hemisphere at 6 h (n=2), 9 h (n=5), and 12 h (n=3) following LPS/PT (**e**).

### Meningeal-route infiltrating neutrophils adopt a distinct transcriptional program enriched for IL-36γ

We next hypothesized that neutrophils entering the brain via the meningeal route may acquire distinct effector programs compared with neutrophils entering from the circulation through vascular routes. To distinguish blood-borne versus meningeal-route cells, we performed dual in vivo labeling using intravenous (IV) anti-CD45-AF488 to tag circulating leukocytes and intracisterna magna (ICM) anti-CD45.2-AF700 to label cells accessible from leptomeningeal spaces (Fig. 6a). Animals were sacrificed 1 h after labeling; blood, meninges, and ipsilateral hemispheres were harvested and analyzed by flow cytometry. Within Live^+^CD11b^+^Ly6G^+^ neutrophils, we identified IV+ (CD45^+^) and ICM+ (CD45.2^+^) populations in meninges and brain at 24 h post-LPS/PT (Fig. 6a). We then sorted five neutrophil populations for bulk RNA sequencing: (1) IV+ neutrophils in blood (Blood), (2) IV+ neutrophils in meninges (IV-MN), (3) ICM+ neutrophils in meninges (ICM-MN), (4) IV+ neutrophils in brain (IV-Brain), and (5) ICM+ neutrophils in brain (ICM-Brain). Principal component analysis revealed that blood neutrophils were transcriptionally distinct from tissue-associated neutrophils in meninges and brain (Fig. 6b). Notably, IV-MN and ICM-MN displayed broadly similar profiles, suggesting that meningeal localization itself strongly shapes neutrophil states. While IV-Brain clustered closely with IV-MN, ICM-Brain diverged substantially from ICM-MN, indicating that neutrophils entering brain from meningeal spaces undergo pronounced transcriptional remodeling upon parenchymal entry (Fig. 6b).

**Figure 6.**
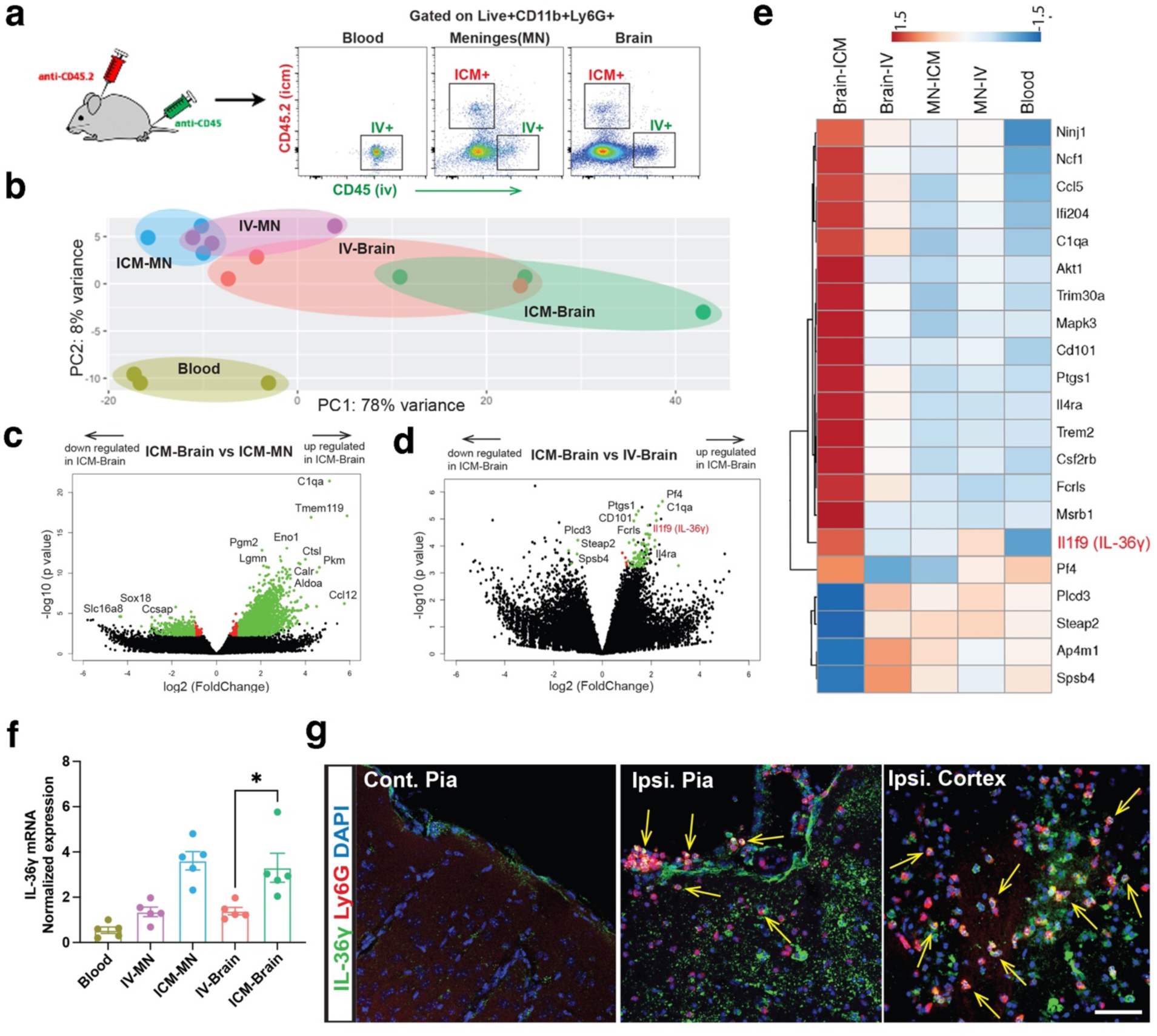
Meningeal neutrophils exhibit elevated IL-36γ expression following LPS-sensitized pediatric stroke. **a,** Animals were euthanized 24 h post-LPS/PT. Neutrophils from distinct anatomical compartments were differentially labeled using intracisternal (ICM, n=3) or intravenous (IV, n=3) CD45.2 antibody. Five neutrophil populations were sorted from blood, meninges (MN), and brain parenchyma for bulk RNA sequencing analysis: Blood, ICM-MN, IV-MN, ICM-Brain, IV-Brain. **b,** Principal component analysis (PCA) of the five neutrophil populations identified by compartment and labeling strategy. **c,** Volcano plot comparing transcriptional profiles of ICM-labeled neutrophils isolated from the brain parenchyma (ICM-Brain) versus meninges (ICM-MN). Genes with an absolute fold change ≥2 or ≤0.5 and meeting statistical significance (adjusted p-value) are highlighted in green, while genes reaching statistical significance with smaller fold changes are shown in red. **d,** Volcano plot comparing transcriptional profiles of ICM-labeled (ICM-Brain) versus IV-labeled (IV-Brain) neutrophils isolated from the brain parenchyma. Differentially expressed genes are color-coded as in (**c**). **e,** Heatmap of selected neutrophil activation–associated genes derived from the significantly differentially expressed gene set in (**d**), shown across all five neutrophil populations. **f,** RT-qPCR validation of Il36g mRNA expression across the five neutrophil populations. Data are presented as mean ± S.E.M. (n = 5 mice for each group). Statistical significance was determined by one-way ANOVA with the Tukey post hoc test (* p<0.05). **g,** post-LPS/PT. Yellow arrows indicate IL-36γ–expressing neutrophils localized to the ipsilateral pia mater and brain parenchyma (n=3).

Differential expression analysis between ICM-Brain and ICM-MN identified 2,680 significantly altered genes (Fig. 6c), with gene set enrichment implicating heightened proteasome activity, oxidative phosphorylation, threonine-type peptidase activity, ATPase activity, and cytosolic transport programs in meningeal-route brain-infiltrating neutrophils (Sup. Table 1). By contrast, comparison between ICM-Brain and IV-Brain identified 81 differentially expressed genes (Fig. 6d; Sup. Table 1). A heatmap of activation-associated genes highlighted distinct expression patterns in ICM-Brain neutrophils, including altered abundance of genes linked to neutrophil activation and effector function (e.g., Cd101, Ncf1, Ifi204, C1qa; Fig. 6E). Among these genes, we identified **Il1f9 (IL-36γ)** as a prominent cytokine enriched in ICM-Brain neutrophils relative to IV-Brain neutrophils (Fig. 6e), suggesting that the meningeal route is associated with induction of a barrier- and inflammation-linked cytokine program. To validate these findings, we quantified Il1f9 (IL-36γ) mRNA by qPCR across the five sorted neutrophil populations. Consistent with bulk RNA-seq, ICM-Brain neutrophils exhibited significantly higher IL-36γ transcripts compared with IV-Brain neutrophils (Fig. 6f). We further confirmed IL-36γ protein expression by immunohistochemistry at 24 h post-LPS/PT, observing IL-36γ+ Ly6G+ neutrophils both along the ipsilateral pial surface and within the ipsilateral cortex (Fig. 6g, yellow arrows). These results identify IL-36γ as a hallmark cytokine associated with meningeal-route brain-infiltrating neutrophils during pediatric stroke.

### Single-cell profiling of meningeal immune cells identifies neutrophils as the dominant IL-36γ source after LPS/PT

To define the cellular source and context of IL-36γ induction in meninges, we performed single-cell RNA sequencing on sorted Live^+^CD45^+^ meningeal immune cells from sham and LPS/PT animals at 24 h post-injury. Across 15,574 cells, unsupervised clustering identified 20 clusters. Using canonical lineage markers, we annotated neutrophil clusters (Csf3r, Camp, Lcn2, Ltf, Mmp9, S100a8/a9), B cell clusters (Ighm, Igkc, Cd79a/b, Pax5), and monocyte/macrophage clusters (Mafb, Saa3, Ccr2, Arg1) (Sup. Table 2; Fig. 7a). In sham meninges, diverse immune populations were present, with B cells comprising a major fraction. In contrast, LPS/PT meninges were dominated by neutrophils (Fig. 7b), consistent with our flow cytometric findings. Within the neutrophil compartment, clusters enriched in LPS/PT displayed increased expression of inflammatory effector molecules (e.g., Il1a, Il1b, Tnf, Icam1) compared with neutrophil clusters predominant in sham animals (data not shown), indicating a distinct activated meningeal neutrophil state induced by LPS-sensitized stroke. Importantly, Il1f9 (IL-36γ) expression localized primarily to neutrophil clusters in LPS/PT meninges (Fig. 7c), independently corroborating our bulk RNA-seq and validation data and supporting neutrophils as a major source of IL-36γ at the meningeal interface in this model. We also detected low-level Il1f9 expression in neutrophil clusters from sham meninges (Fig. 7c), raising the possibility that IL-36γ may participate in basal meningeal immune–barrier homeostasis, with strong induction occurring under inflammatory priming and injury.

**Figure 7.**
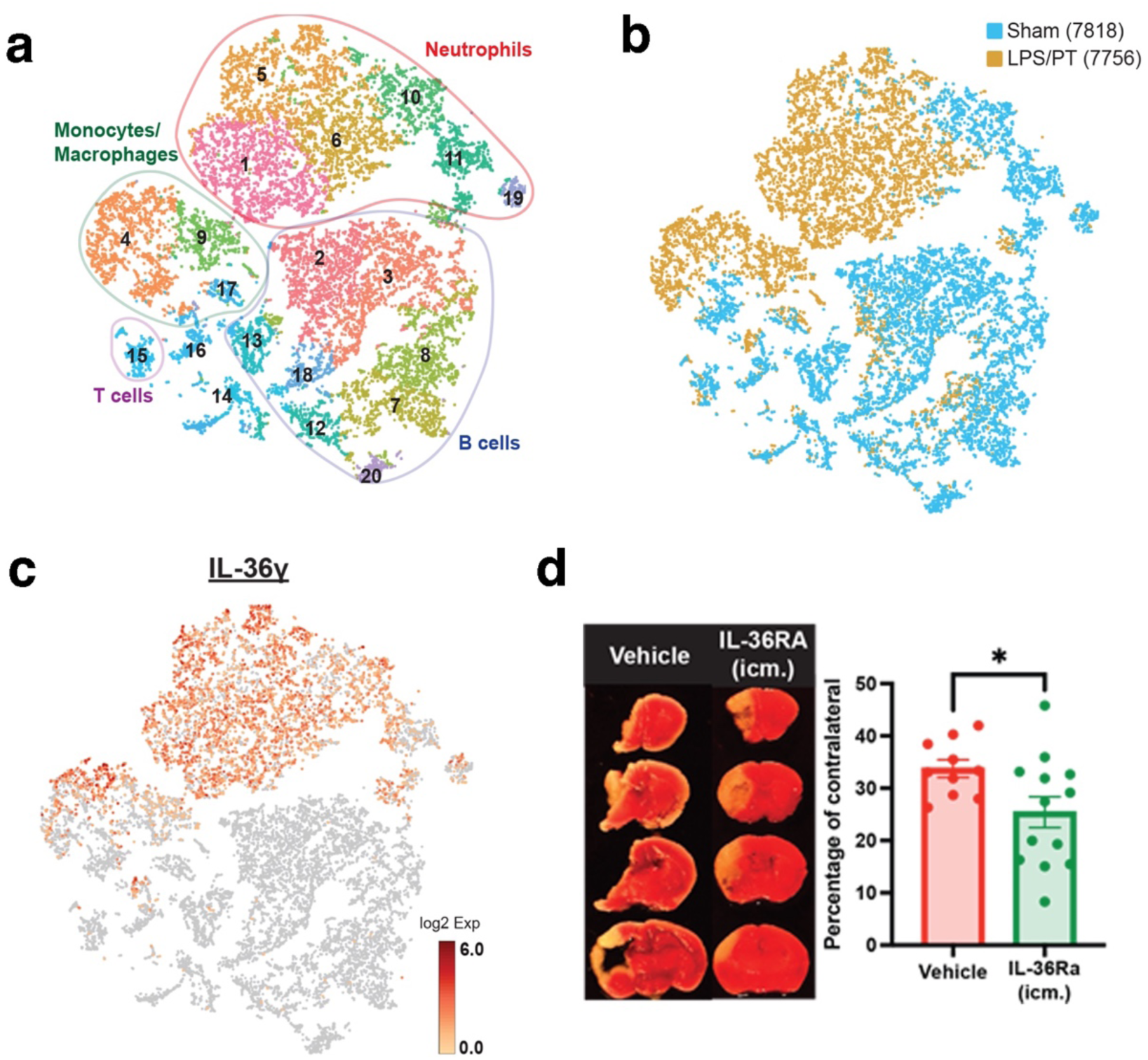
Single-cell RNA sequencing of meningeal immune cells identifies neutrophils as a major source of IL-36γ, which exacerbates infarct severity following LPS-sensitized pediatric stroke. **a,** Single-cell RNA sequencing was performed using the 10x Genomics platform on CD45⁺ immune cells isolated from the meninges of sham-operated and LPS-sensitized stroke animals at 24 h post-stroke. Major immune cell populations were identified by unsupervised clustering and annotated based on canonical marker gene expression. **b,** t-distributed stochastic neighbor embedding (t-SNE) visualization of meningeal immune cells from sham (7,818 cells) and LPS/PT (7,756 cells) conditions. **c,** Feature plot showing Il36g expression across distinct immune cell clusters in the meningeal single-cell RNA-sequencing dataset. **d,** Mice received intracisternal administration of IL-36 receptor antagonist (IL-36RA) at 3 h post-LPS/PT (n=13). Infarct size were quantified by TTC staining at 3 days post-stroke and expressed as a percentage of the contralateral hemisphere. Data are presented as mean ± S.E.M. * p < 0.05 compared with the vehicle-treated group (n = 8) by unpaired t-test.

### Local inhibition of IL-36 signaling reduces infarct size, supporting a detrimental role for the meningeal neutrophil–IL-36γ axis

Given the enrichment of IL-36γ in meningeal-route infiltrating neutrophils and the established role of IL-36 family cytokines in barrier inflammation across epithelial tissues, we tested whether IL-36 signaling contributes functionally to pediatric stroke pathology in LPS/PT. We administered recombinant IL-36 receptor antagonist (IL-36Ra/IL-1F5) via intracisterna magna injection at 3 h post-LPS/PT and quantified infarct volume by TTC staining at day 3 (Fig. 7d). IL-36Ra treatment significantly reduced infarct size compared with isotype-treated controls (Fig. 7d), supporting a detrimental role for IL-36 signaling in inflammation-sensitized pediatric stroke.

### Local blockade of ICAM-1 reduces neutrophil entry into parenchyma and improves outcome in LPS/PT

Neutrophil adhesion and transmigration commonly rely on ICAM-1/LFA-1 interactions^31^. To test whether this pathway contributes to neutrophil movement across the brain–meningeal interface, we delivered anti-ICAM-1 antibody via intracisterna magna injection at 3 h post-LPS/PT and quantified neutrophils in the parenchyma at 24 h. Compared with isotype-treated controls, ICM administration of anti-ICAM-1 markedly reduced parenchymal neutrophil accumulation (Fig. 8a–b). In contrast, meningeal neutrophil numbers were not significantly altered (Fig. 8c), indicating that ICAM-1 blockade primarily inhibits the migration step from meningeal compartments into parenchyma rather than initial recruitment into meninges. Because systemic anti-ICAM-1 therapy (enlimomab) failed clinically and was associated with increased adverse events^32^, we next asked whether localized blockade confined to meningeal spaces could provide benefit without broadly suppressing systemic immune function. Local (ICM) anti-ICAM-1 treatment reduced pro-inflammatory cytokine transcripts (Tnf, Il6) in the ipsilateral hemisphere (Fig. 8d) and significantly decreased infarct volume at day 3 post-LPS/PT (Fig. 8e). By contrast, systemic anti-ICAM-1 administration (intraperitoneal) did not significantly reduce infarct size compared with isotype controls (Fig. 8f). Collectively, these results suggest that neutrophil migration across the brain–meningeal interface is an ICAM-1–dependent step that is therapeutically targetable via localized meningeal delivery.

**Figure 8.**
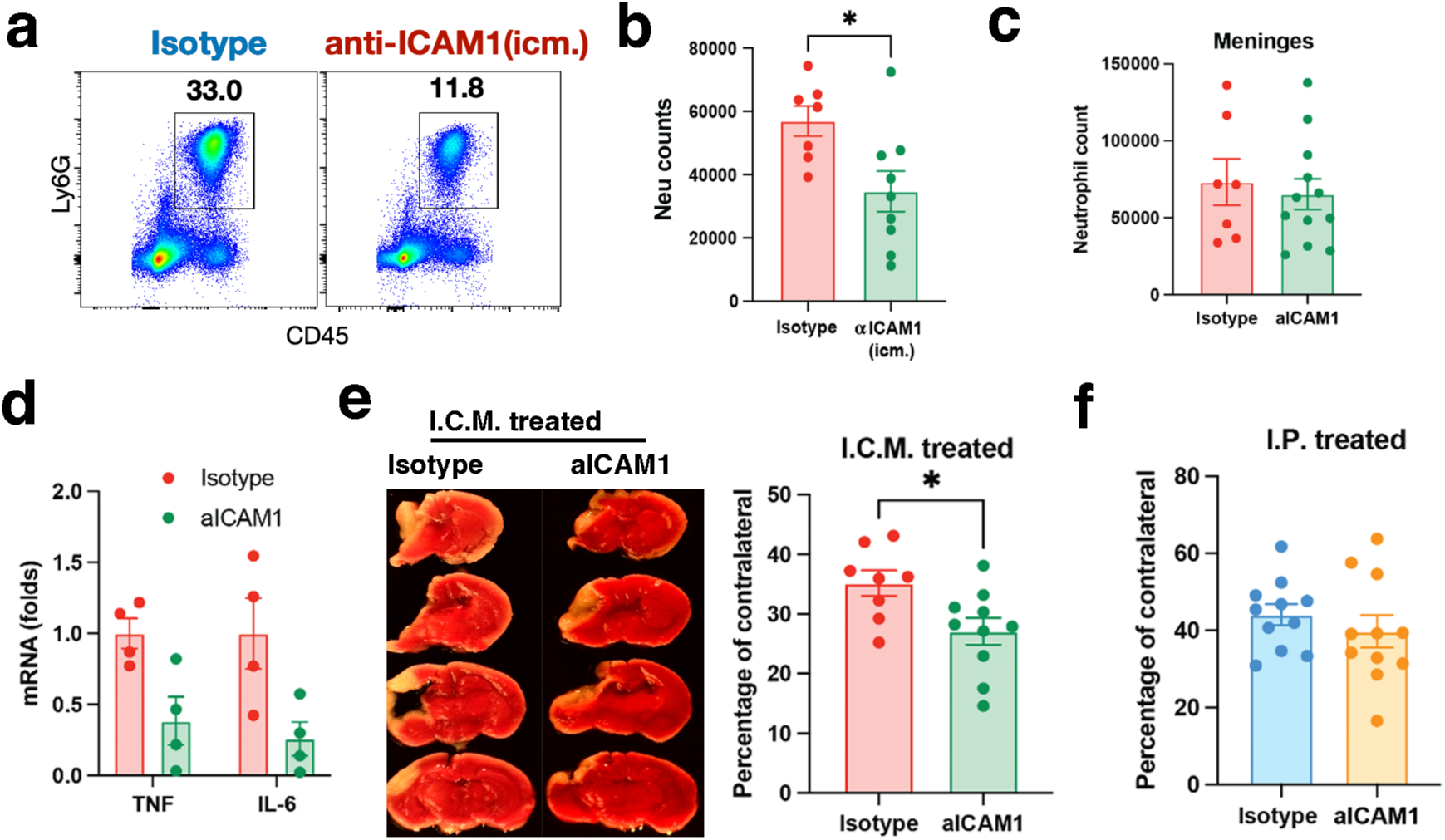
Blockade of ICAM-1 in the leptomeningeal space reduces neutrophil migration into the brain parenchyma and attenuates infarct size in LPS-sensitized pediatric stroke. **a-c,** LPS/PT animals received an intracisternal injection of an ICAM-1–blocking antibody (5 mg/mL; Bio X Cell; 2 µL) at 3 h post-stroke. Representative flow cytometric plots from the ipsilateral brain hemisphere at 24 h post-stroke are shown (**a**). Quantification of neutrophils (gated as CD45hiCD11bhiLy6G⁺) in the ipsilateral brain hemisphere at 24 h post-stroke following intracisternal ICAM-1 blockade (n=9) or isotype control treatment (n=7) (**b**). Quantification of neutrophils (CD45hiCD11bhiLy6G⁺) in the meninges (dura mater) at 24 h post-stroke following intracisternal ICAM-1 blockade (n=12) or isotype control treatment (n=7). Data are presented as mean ± S.E.M. p < 0.05 compared with the isotype group by unpaired t-test (**c**). **d,** Animals from each experimental group were euthanized 24 h post-stroke, and ipsilateral brain hemispheres were collected for quantitative PCR analysis of Tnf and Il6 mRNA expression (n=4 for each group). **e,** Mice treated intracisternally with anti–ICAM-1 antibody (n=10) or isotype control (n=8) were euthanized 3 days post-stroke. Brain sections were stained with TTC, infarct size were quantified, and representative TTC-stained sections are shown. **f,** Animals treated systemically (intraperitoneal injection) with anti–ICAM-1 antibody or isotype control were euthanized 3 days post-stroke. Brain sections were stained with TTC, and infarct size were quantified (n=11 per group). *P < 0.05, compared with the Isotype group by unpaired t-test.

**Figure 9.**
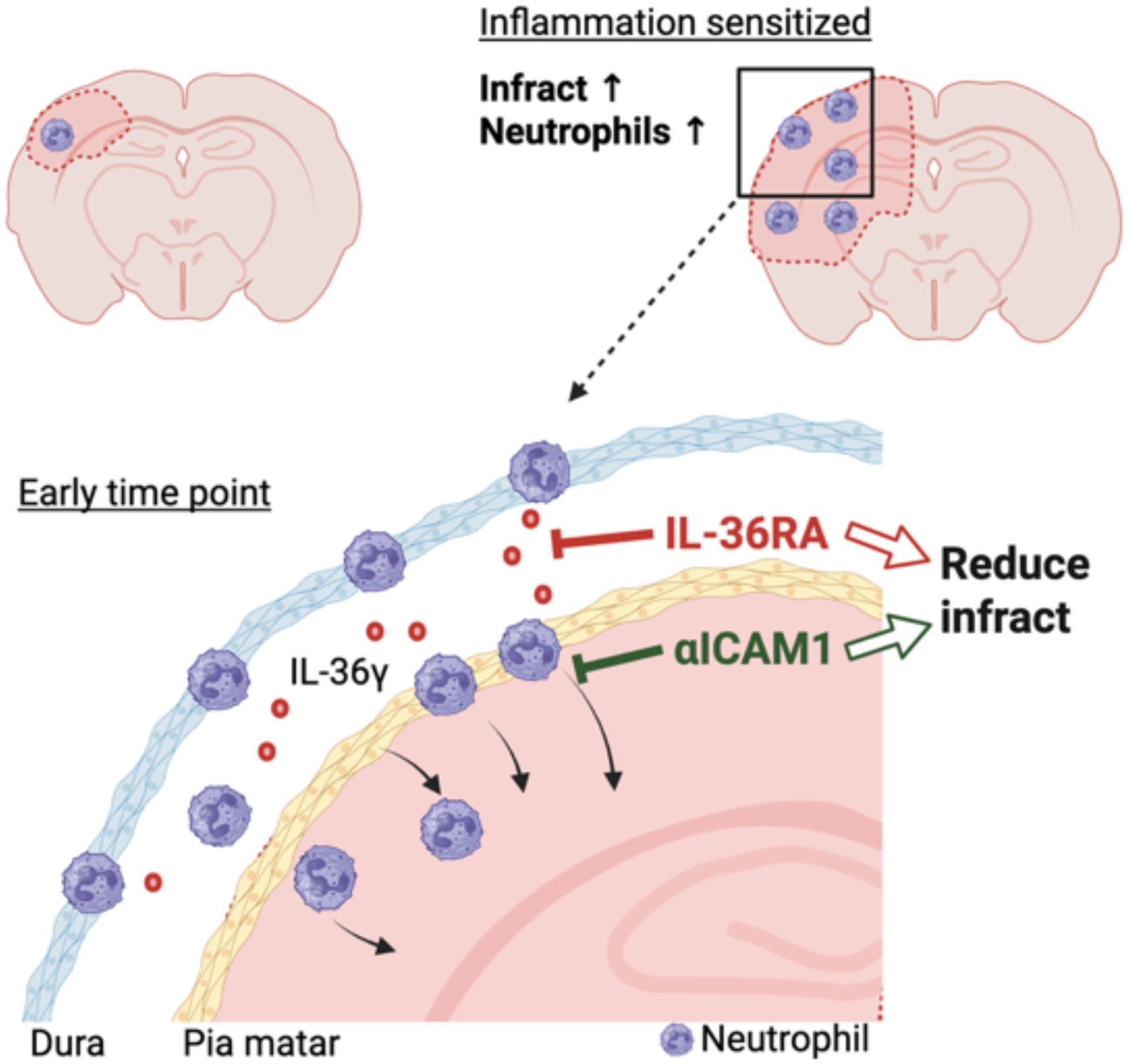
Schematic summary of leptomeningeal neutrophil–IL-36γ signaling in LPS-sensitized pediatric stroke. LPS sensitization exacerbates infarct severity following pediatric stroke by amplifying neutrophil-driven inflammation. In the early phase, neutrophils accumulate in the leptomeninges and subsequently migrate across the pia mater into the cortical parenchyma. Meningeal neutrophils exhibit elevated IL-36γ expression, which promotes inflammatory amplification and tissue injury. Disruption of the brain–meningeal barrier facilitates trans-pial neutrophil trafficking. Intracisternal administration of IL-36 receptor antagonist (IL-36RA) or blockade of ICAM-1 within the leptomeningeal space reduces neutrophil infiltration and attenuates infarct size. These findings identify the leptomeninges as a critical inflammatory gateway and highlight IL-36γ signaling and ICAM-1–mediated neutrophil migration as therapeutic targets in inflammation-sensitized pediatric stroke.

## Discussion

In this study, we developed an inflammation-sensitized pediatric stroke model and examined how innate immune infiltrates contribute to worsened ischemic injury in the immature brain. Using low-dose LPS priming prior to photothrombotic stroke at P16, we found markedly increased infarct burden accompanied by a dominant influx of neutrophils with a pro-inflammatory activation profile. Functional experiments supported a central role for neutrophils in this setting: antibody-mediated neutrophil depletion reduced infarct volume in LPS/PT animals, whereas genetic disruption of CCR2-dependent monocyte recruitment did not provide protection. Together, these findings indicate that inflammatory priming shifts the acute cellular contributors to pediatric ischemic injury toward a neutrophil-driven response. A notable contrast in our dataset is the minimal impact of CCR2-dependent monocyte recruitment in LPS/PT, compared with our prior work in neonatal hypoxic–ischemic (H-I) injury where monocytes contributed to pathology and could adopt microglia-like programs^10^. Several factors may contribute to this difference. First, distinct CNS injury paradigms may preferentially engage different entry portals and compartments. In our LPS/PT model, we observed prominent leukocyte accumulation at the pial/leptomeningeal interface, whereas we did not detect enhanced immune accumulation at the choroid plexus^33, 34^ under these conditions (data not shown). Second, LPS priming produced a cytokine/chemokine milieu enriched for neutrophil-attracting signals (e.g., CXCL1 and CXCL2), which likely favors rapid neutrophil recruitment and may diminish the relative contribution of monocyte-mediated mechanisms during the acute window. Third, developmental maturation may shape the effector capacity of myeloid subsets. Neonatal neutrophils are often described as functionally distinct from adult neutrophils, with reduced canonical effector outputs in certain contexts. By P16, neutrophils may be more functionally mature, potentially increasing their capacity for adhesion, transmigration, and inflammatory effector responses^8, 35^. Although these considerations are consistent with our observations, defining the precise age-dependent mechanisms will require direct comparisons across developmental stages and matched injury paradigms.

Beyond identifying neutrophils as key contributors, our results highlight meningeal compartments as an early site of neutrophil accumulation and a likely route of entry into injured parenchyma. In adult stroke models, BBB disruption and endothelial activation can occur early after ischemia^29^, yet in our model we observed minimal neutrophil presence in the parenchyma at 6h-12h post-injury despite early induction of neutrophil chemoattractants. In contrast, neutrophils accumulated in meninges as early as 6 h and remained elevated through 24h. Histological mapping relative to laminin and GFAP further indicated that neutrophils localize on the CSF-facing side of the pia early, then appear to traverse the pia and glia limitans around 12 h (Fig 3e), coinciding with reduced laminin signal and disruption of the glia limitans at later time points. Consistent with these spatial observations, tracer-based assays demonstrated increased leakage of CSF-Dextran dye by ∼12 h post-LPS/PT, suggesting that functional impairment of the brain–meningeal barrier develops during the window when neutrophils begin to cross the brain surface. Importantly, we directly visualized this meningeal route using complementary approaches. Two-photon imaging captured neutrophils positioned along the pial surface and entering the parenchyma, and KikGR photoconversion provided tracking evidence that a fraction of early brain-infiltrating neutrophils originated from photoconverted meningeal compartments rather than the circulating pool. While our experiments do not definitively distinguish whether early meningeal neutrophils arise primarily from skull bone marrow^36^, meningeal vasculature, or systemic circulation, our IV/ICM labeling strategy supports the presence of neutrophil populations that are accessible from leptomeningeal spaces and behave differently from classically blood-borne neutrophils. These findings are consistent with an emerging view that the meninges are not only an immune reservoir but also a dynamic interface that can shape immune cell trafficking into the CNS during inflammation. Mast cells in meninges have been recently reported guard the gateway of immune cell infiltration and CSF regulation after CNS injury^37, 38^. In adult stroke models, neutrophil accumulation has also been reported in leptomeningeal and perivascular locations after ischemia^39^, suggesting that surface and perivascular compartments may serve as common staging sites for neutrophils, although their relative importance may be heightened in pediatric settings where classical BBB leakage and early parenchymal infiltration can be reduced.

A second major finding is that neutrophils entering the brain via meningeal-accessible routes display a distinct transcriptional program relative to blood-borne brain-infiltrating neutrophils. Bulk RNA sequencing of neutrophil populations defined by IV versus ICM labeling revealed route-associated differences, including enrichment of activation-related genes and the prominent induction of Il1f9 (IL-36γ) in meningeal-route brain-infiltrating neutrophils. We validated this observation by qPCR and immunostaining, and independently confirmed IL-36γ enrichment within meningeal neutrophil clusters by single-cell RNA sequencing. Together, these orthogonal datasets support IL-36γ as a reproducible feature of the meningeal neutrophil response in inflammation-sensitized pediatric stroke. The IL-36 cytokine family has emerged as an important regulator of inflammatory responses at barrier interfaces. IL-36 require proteolytic processing to achieve full biological activity and can amplify local inflammatory programs by inducing chemokines that recruit neutrophils and other immune cells^19^. Neutrophils themselves are a major source of the serine proteases capable of activating IL-36 cytokines^20^, suggesting that neutrophil accumulation at barrier sites may locally enhance IL-36 signaling through a feed-forward inflammatory loop. Consistent with this concept, previous work in experimental autoimmune encephalomyelitis (EAE) demonstrated that neutrophils can express IL-36γ^40^ and that glial cells respond to IL-36 stimulation by producing cytokines and chemokines that further recruit neutrophils to the inflamed CNS environment^40^. In our pediatric stroke model, we similarly observed robust IL-36γ expression in meningeal-associated neutrophils and confirmed IL-36γ protein localization at the pial surface and within the ipsilateral cortex. Importantly, we also detected IL-36 receptor expression on both GFAP⁺ astrocytes and IBA1⁺ microglia in the injured hemisphere (data not shown), suggesting that CNS-resident glial cells are potential responders to neutrophil-derived IL-36γ in this context. Given the known capacity of astrocytes and microglia to produce inflammatory mediators and chemokines, IL-36 signaling in these cells may contribute to amplification of local neuroinflammation and sustained recruitment of innate immune cells after stroke. In addition to glial cells, IL-36R has been reported in epithelial and stromal barrier cells in other tissues, where IL-36 signaling activates NF-κB and MAPK pathways and modulates barrier integrity and inflammatory responses^41^. Taken together, these findings support a model in which IL-36γ produced by meningeal-associated neutrophils engages IL-36R on CNS-resident cells to amplify inflammatory signaling at the brain–meningeal interface. Consistent with this interpretation, pharmacological blockade of IL-36 signaling through intracisternal administration of IL-36Ra significantly reduced infarct size in our model, indicating that IL-36 signaling contributes functionally to inflammation-driven injury in pediatric stroke.

Our data also suggest a locally targetable therapeutic opportunity at meningeal interfaces. Neutrophil transmigration is classically regulated by adhesion pathways including ICAM-1/LFA-1 interactions. We found that intracisternal anti-ICAM-1 reduced parenchymal neutrophil accumulation without reducing meningeal neutrophil numbers, indicating that ICAM-1 blockade primarily affects the step of migration across brain surface barriers rather than recruitment into meninges. Notably, systemic anti-ICAM-1 therapy failed in clinical stroke trials and was associated with adverse outcomes, likely reflecting broad systemic immune perturbation^32^. In contrast, the localized efficacy of intracisternal anti-ICAM-1 in our model raises the possibility that spatially restricted blockade at meningeal interfaces may mitigate damaging neuroinflammation while avoiding systemic immunosuppression. While translation of intracisternal delivery approaches requires substantial additional validation, these results help define the meningeal compartment as a potential therapeutic window in inflammation-sensitized pediatric stroke.

This study has several limitations. First, our barrier permeability assays used macromolecular tracers (e.g., BSA and ∼70kD dextrans), which may underestimate earlier or more subtle permeability changes detectable with smaller tracers. Prior studies have reported very early BBB permeability changes in transient ischemia/reperfusion models using smaller tracers (3kD) **[29]**. A systematic analysis using a range of tracer sizes will be important to define the temporal dynamics and size selectivity of brain–meningeal barrier permeability after pediatric stroke. Second, photothrombotic stroke involves laser illumination, which could potentially influence meningeal inflammation. Although our controls suggest limited immune recruitment attributable to laser exposure alone (data not shown), validation in different ischemia models will strengthen our findings.

In summary, our findings support a framework in which systemic inflammatory priming exacerbates pediatric ischemic injury through a neutrophil-dominant response that is organized early at meningeal interfaces. We identify functional impairment of the brain–meningeal barrier as a route that enables neutrophil entry into the injured immature brain and define a route-associated neutrophil program enriched for IL-36γ. The protective effects of intracisternal IL-36Ra and anti-ICAM-1 suggest that targeting meningeal neutrophil function or transmigration may represent a promising strategy to mitigate inflammation-driven pediatric stroke pathology.

---

**Supplement Figure 1:**
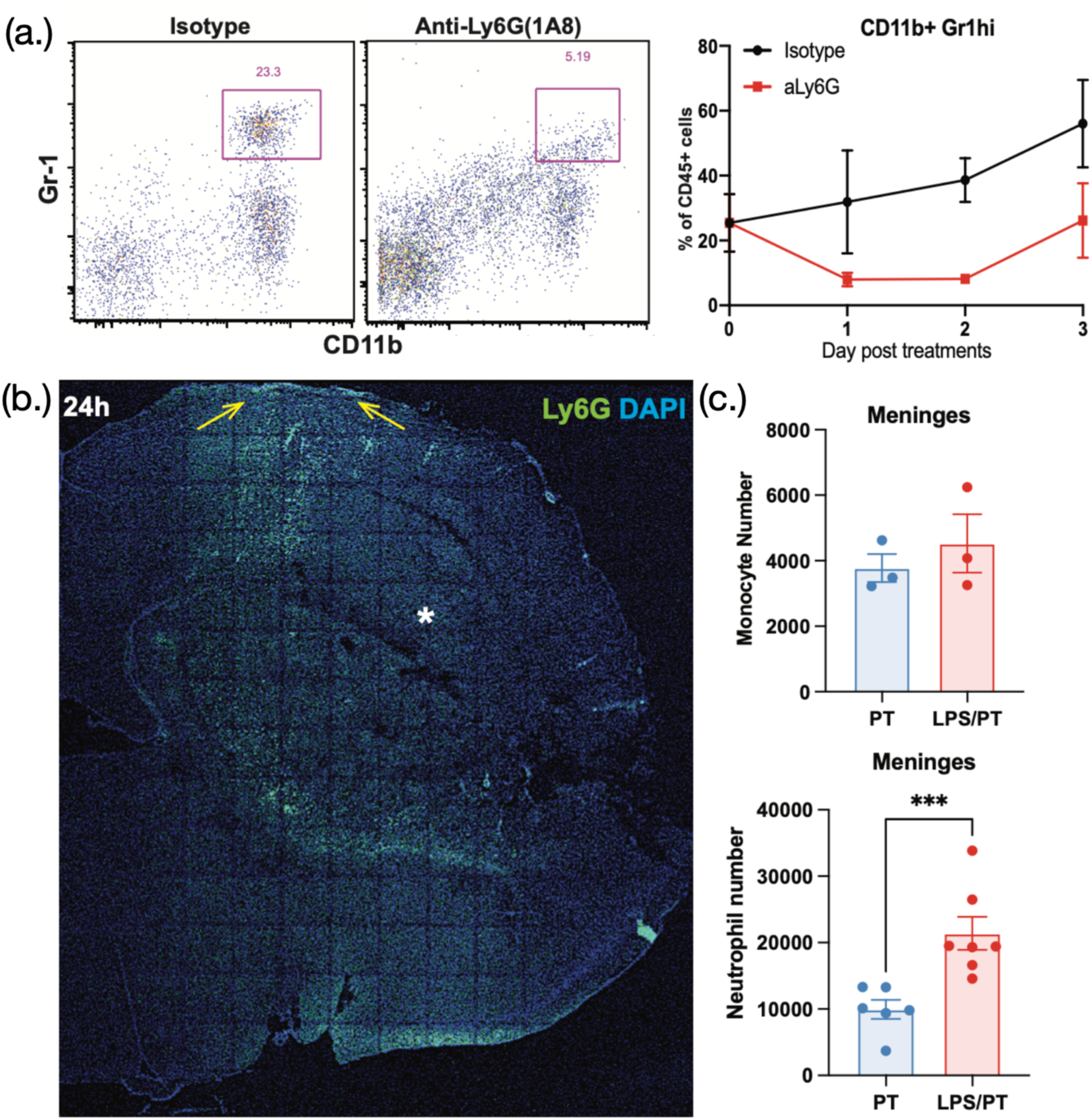
Fig. 1a: Representative flow cytometry plots demonstrating neutrophil depletion following anti-Ly6G (1A8) antibody treatment. Neutrophils were defined as CD11b⁺Gr1^hi^ cells in this experimental setting. The summary graph quantifies the frequency of CD11b⁺Gr1^hi^ neutrophils within total CD45⁺ cells in isotype control– and anti-Ly6G (1A8)–treated animals (n = 4). Fig. 1b: Tile-scan image of the ipsilateral hemisphere at 24 h following LPS/PT. Sections were stained for Ly6G (green) and DAPI (blue), and the infarct region is indicated by a star. Notably, neutrophils accumulated along the pia mater surrounding the infarct area, as indicated by yellow arrows. Fig. 1c: Quantification of meningeal monocytes and neutrophils in PT and LPS/PT groups at 24 h post-stroke.

